# Large structural variants in KOLF2.1J are unlikely to compromise neurological disease modelling

**DOI:** 10.1101/2024.01.29.577739

**Authors:** Mallory Ryan, Justin A. McDonough, Michael E. Ward, Mark R. Cookson, William C. Skarnes, Florian T. Merkle

## Abstract

Gracia-Diaz and colleagues analysed high-density DNA microarray and whole genome sequencing (WGS) data from the KOLF2.1J ‘reference’ human induced pluripotent stem cell (hiPSC) line^1^, and report the presence of five high-confidence heterozygous copy number variants (CNVs) at least 100kbp in length^2^. Since three of these CNVs span coding genes, some of which have been associated with neurodevelopmental disease, the authors raise the concern that these CNVs may compromise the utility of KOLF2.1J for neurological disease modelling. We appreciate their thorough analysis and thoughtful interpretation, and agree that potential users of this line should be made aware of all cases where KOLF2.1J differs from the reference genome. However, we believe that the benefits from the widespread use of KOLF2.1J outweigh the potential risks that might arise from the identified CNVs.

## Results

To corroborate the findings of Gracia-Diaz and colleagues, we performed llumina Infinium Global Diversity Array with Cytogenetics-8 DNA microarray analysis on KOLF2.1J and parental KOLF2_C1 cells, a clonal subline of the original KOLF2 parental cell line (HPSI0114i-kolf_2) used to generate KOLF2.1J. We confirmed the presence of the five reported CNVs in both KOLF2.1J and KOLF2-C1 (Fig. S1A,B). Furthermore, we have not observed other CNVs arising at appreciable frequencies in our extensive CRISPR/Cas9-based editing of KOLF2.1J. We believe that the most parsimonious explanation is that the five CNVs were already present in a subpopulation of the original KOLF2 parental cell line or of the fibroblasts used for reprogramming. Mosaicism in parental iPS cell lines is common and difficult to detect on SNP arrays, highlighting the importance of isolating sublines prior to deep characterization and editing of a reference cell line, as we performed for KOLF2.1J.

Germline CNVs larger than 100kbp are individually rare but collectively common, affecting a majority (∼65-80%) of individuals in human populations^3^. In other words, there are no truly ‘wild-type’ individuals, and most of them will present large CNVs that deviate from reference genomes. While some CNVs contribute to disease, the contribution of a given CNV is often difficult to discern. The reported CNVs in KOLF2.1J affect the genes *JARID2* and *ASTN2*, which are both predicted to be dosage-sensitive in recent studies predicting genome-wide dosage sensitivity and gene mutational constraint^4,5^ (Fig. S1C). We therefore discuss these genes in greater detail below.

Although large deletions and frameshift mutations affecting *JARID2* have been associated with a human neurodevelopmental disorder, the majority of established pathogenic CNVs also encompass other genes on chr6p22^6^, *Jarid2* heterozygous mice do not have reported developmental phenotypes^7^, and *JARID2* is not annotated as dosage sensitive in ClinGen (https://search.clinicalgenome.org/kb/gene-dosage) arguing against penetrant, evolutionarily conserved haploinsufficiency. Furthermore, many groups have shown that KOLF2.1J differentiates normally to diverse cell types, particularly neural lineages. The transcriptomes of these KOLF2.1J-derived cells also resemble those derived from other iPSC cell lines, suggesting that any potential *JARID2* gene dosage effects on cellular differentiation are difficult to detect using current iPSC differentiation methods.

*ASTN2* encodes a brain-expressed protein involved in synapse dynamics whose deletion is associated with schizophrenia, although again most cases involve multiple gene disruptions on chr9q33.1. No obvious neurological phenotypes have been reported in mice heterozygous for a deletion encompassing *Astn2* in the absence of other gene deletions^8^, nor is *ASTN2* annotated as dosage sensitive. We note that our previous data showed that KOLF2.1J-derived neurons appeared to support synaptic activity^1^.

Overall, we agree that the findings of Diaz-Garcia and colleagues are robust and that it is important for the community to understand that KOLF2.1J likely has only one functional allele of *JARID2* and *ASTN2*. We suggest that groups with particular interest in studying the biology of either of these genes may wish to select another cell line for their studies. However, there are several reasons to believe that KOLF2.1J remains an attractive choice of iPSC line for most purposes.

First, the functional consequences of deletions at chr6p22 and chr9q33 in human iPSCs remain unclear, and the apparently equivalent differentiation potential of KOLF2.1J compared to other commonly used lines argues that KOLF2.1J is functionally useful. Second, KOLF2.1J has already been widely distributed to groups around the world by The Jackson Laboratory (https://www.jax.org/jax-mice-and-services/ipsc), and forms the basis of several large-scale initiatives including iNDI focusing on highly penetrant variants associated with neurodegeneration, MorPhiC (https://morphic.bio/) focusing on cellular phenotypes arising from gene knockout, and SSPsyGene (https://sspsygene.ucsc.edu/) focusing on neurodevelopmental disorders. The KOLF2.1J iPSC line is readily available for research and commercial use, and the deep phenotyping and the wealth of data generated from it and its derivatives across diverse genotypes will be very useful to the community. Third, to our knowledge other iPSC lines have not been as deeply scrutinised as KOLF2.1J, and we expect that other lines are likely to show other genetic variants of unclear functional relevance upon closer scrutiny, especially as new technologies are brought to bear. For example, a recent comparison of inbred mouse strains using long-read sequencing revealed the presence of approximately 5 structural variants in every gene across the mouse genome, including hundreds of variants per strain that are predicted to ablate gene function^9^. These findings do not invalidate the utility of using inbred mouse strains or a given cell line, but highlight the fact that the more we understand about their genetic backgrounds, the more confidently we can interpret resulting phenotypes. However, we acknowledge that it is best practice to replicate any key mutation- or knockout-associated phenotypes observed in KOLF2.1J in multiple additional lines to confirm they are robust to genetic background. Indeed, we are currently extending the iNDI project to include additional background lines. We are therefore grateful to Gracia-Diaz and colleagues for bringing the variants in KOLF2.1J to the attention of the community.

## Acknowledgements

F.T.M. is a New York Stem Cell Foundation - Robertson Investigator (NYSCF-R-156) and is supported by the Wellcome Trust and Royal Society (211221/Z/18/Z) and a Ben Barres Early Career Acceleration Award from the Chan Zuckerberg Initiative’s Neurodegeneration Challenge Network (CZI NDCN 191942). This research was supported in part by the Intramural Research Program of the NIH, National Institute on Aging (NIA), National Institutes of Health, Department of Health and Human Services; project number [ZO1 AG000535], as well as the National Institute of Neurological Disorders and Stroke. We gratefully acknowledge the contribution of the Scientific Services at the Jackson Laboratory. For the purpose of open access, the authors have applied a CC-BY public copyright license to any Author Accepted Manuscript version arising from this submission.

## Author contribution

F.T.M. wrote the letter, with input from all co-authors. M.R. analysed DNA microarray data under the supervision of J.M. and W.C.S.

## Declaration of Interests

The authors declare no competing interests.

---

**Figure S1.**
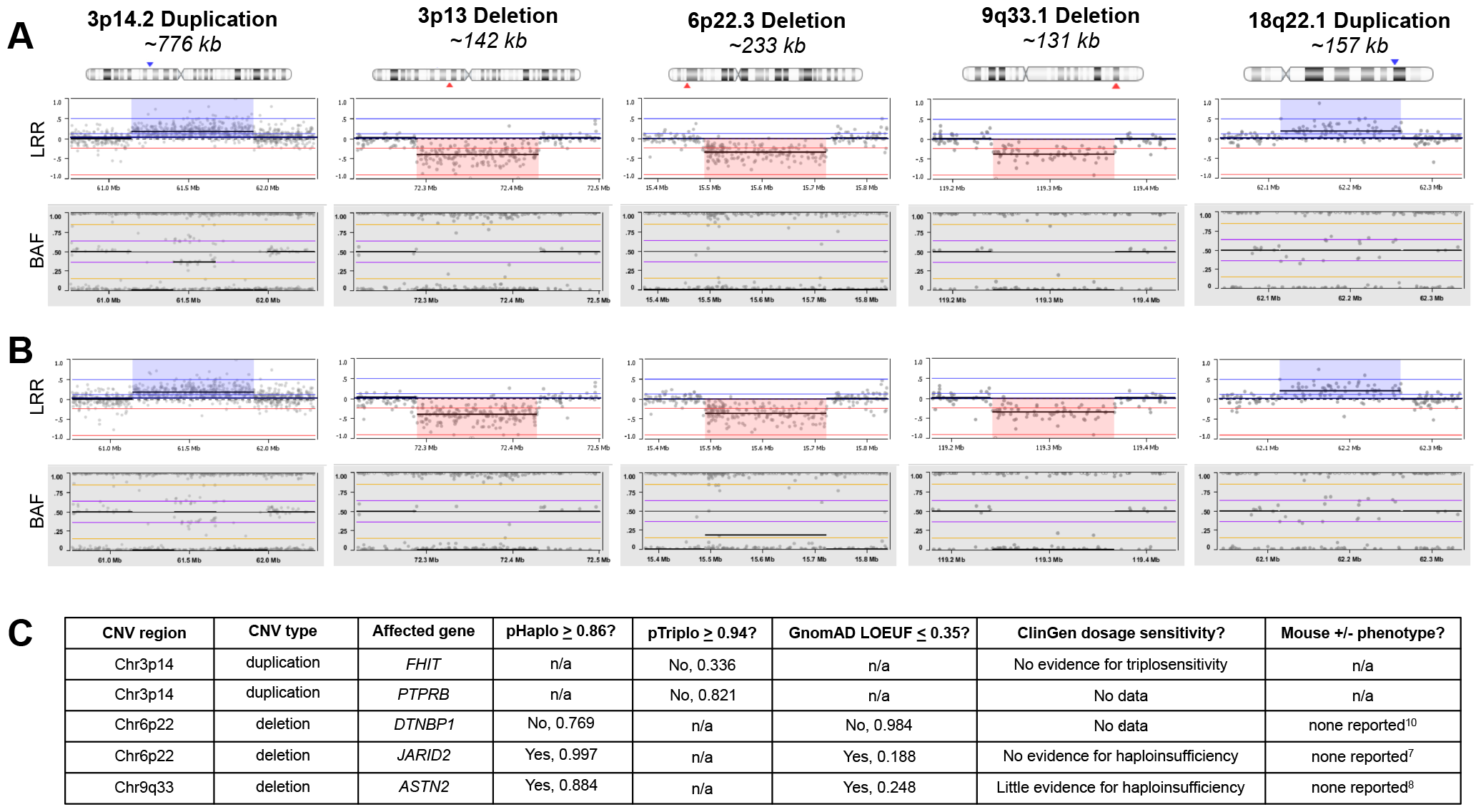
Copy number variants in KOLF2.1J and parental KOLF2_C1 iPSC lines. **A**,**B)** Plots of structural variants from DNA microarray data called by Log R Ratio (LRR) and B Allele Frequency (BAF) in KOLF2.1J (A) and KOLF2_C1 (B) showing two duplicated regions and three deleted regions visualised by VIA software v7.0 (Bionano Genomics). **C)** While bioinformatic tools predict potential pathogenicity of heterozygous variants in Chr6p22 and Chr9p33 deletions, manual annotation in ClinGen and heterozygous mouse models do not support this interpretation.

## Notes

### Competing Interest Statement

The authors have declared no competing interest.

